# Defining Major Depressive Disorder Cohorts Using the EHR: Multiple Phenotypes Based on ICD-9 Codes and Medication Orders

**DOI:** 10.1101/227561

**Authors:** Wendy Marie Ingram, Anna M. Baker, Christopher R. Bauer, Jason P. Brown, Fernando S. Goes, Sharon Larson, Peter P. Zandi

**Affiliations:** Department of Mental Health, Johns Hopkins Bloomberg School of Public Health, Baltimore, Maryland, USA; Department of Psychiatry, Geisinger Health System, Danville, Pennsylvania, USA; Department of Psychology, Bucknell University, Lewisburg, Pennsylvania, USA; Biomedical and Translational Informatics Institute, Geisinger Health System, Danville, Pennsylvania, USA; Department of Psychiatry and Behavioral Sciences, Johns Hopkins School of Medicine, Baltimore, Maryland, USA; College of Population Health, Thomas Jefferson University, Philadelphia, Pennsylvania, USA

**Keywords:** Depression, Electronic health records, Phenotypic algorithms, Clinical informatics

## Abstract

**Background:** Major Depressive Disorder (MDD) is one of the most common mental illnesses and a leading cause of disability worldwide. Electronic Health Records (EHR) allow researchers to conduct unprecedented large-scale observational studies investigating MDD, its disease development and its interaction with other health outcomes. While there exist methods to classify patients as clear cases or controls, given specific data requirements, there are presently no simple, generalizable, and validated methods to classify an entire patient population into varying groups of depression likelihood and severity.

**Methods:** We have tested a simple, pragmatic electronic phenotype algorithm that classifies patients into one of five mutually exclusive, ordinal groups, varying in depression phenotype. Using data from an integrated health system on 278,026 patients from a 10-year study period we have tested the convergent validity of these constructs using measures of external validation, including patterns of psychiatric prescriptions, symptom severity, indicators of suicidality, comorbidity, mortality, health care utilization, and polygenic risk scores for MDD.

**Results:** We found consistent patterns of increasing morbidity and/or adverse outcomes across the five groups, providing evidence for convergent validity.

**Limitations:** The study population is from a single rural integrated health system which is predominantly white, possibly limiting its generalizability.

**Conclusion:** Our study provides initial evidence that a simple algorithm, generalizable to most EHR data sets, provides categories with meaningful face and convergent validity that can be used for stratification of an entire patient population.

## INTRODUCTION

Depression is a highly prevalent mental illness that accounts for $43 billion in medical costs annually and is a leading cause of disability (Maurer, 2012). Depression has been linked to worse outcomes and increased healthcare utilization for numerous common medical disorders (Ang et al., 2005; Celano and Huffman, 2011; Pouwer et al., 2013; Thomas and Price, 2008; Young et al., 2010). However, depression is a heterogeneous disorder, and its etiologies remain poorly understood (Kalia, 2005). There is an urgent need to better understand the causes and course of depression in order to develop more effective treatment and prevention strategies. Electronic Health Records (EHR) from large integrated health systems now offer the opportunity for researchers to conduct unprecedented, large scale studies of patients in real-world settings (Affolter et al., 2017; Fiest et al., 2014; Jin et al., 2015; Morley et al., 2014; Newton-Dame et al., 2016; Simon et al., 2015). Critical to these pursuits are phenotypic algorithms that distinguish who has the disorder within the patient population (McDonald et al., 2016; Sallakh et al., 2017), versus those who may have related disorders, versus those who should be considered unaffected. Depression is a particularly difficult phenotype to define and studies often use heterogeneous criteria when utilizing EHR data to identify patients with depression (Anderson et al., 2015; Huang et al., 2014; Madden et al., 2016; Mayer et al., 2017; Pathak et al., 2014).

As with any phenotypic algorithm, the challenge is to validly define depression with high sensitivity and specificity, limiting both false positive and false negative classification of patients (Cole et al., 2016). An additional complexity is the fact that depression likely exists on a continuum with a range of severity in a population. There are at least four potential sources of information from the EHR for defining depression: a) International Classification of Diseases (ICD) diagnosis codes; b) depression screening measures; c) medication orders; and d) clinical notes. While some studies use only ICD diagnosis codes to identify patients (Gill et al., 2008; Mayer et al., 2017), others have demonstrated that using these alone has inferior sensitivity and precision when compared with combinatorial models that use multiple sources of information (Perlis et al., 2012). National recommendations that adults be screened for depression annually has increased the availability and use of symptom questionnaires (Anderson et al., 2015; Huang et al., 2014; Pathak et al., 2014; U.S. Preventive Services Task Force, 2009) such as the Patient Health Questionnaire (PHQ-9) (Kroenke et al., 2001a). However, implementation of such screening measures has been fairly recent, is not standardized, and shows limited agreement with ICD codes for depression (Kobus et al., 2013; Madden et al., 2016), possibly due to successful treatment of the depression. Phenotyping algorithms may also use information from medication treatment codes (Conway et al., 2011; Nadkarni et al., 2014; Newton et al., 2013). Unfortunately, there may be a long delay between when patients with depression first experience the onset of symptoms and ultimately receive care, including with medication (Dockery et al., 2015; Green et al., 2012; Thompson et al., 2004; Wang et al., 2005). Additionally, antidepressants may be prescribed for a variety of comorbid mental (Shah and Han, 2015) and non-mental health indications (Wong et al., 2017), such as tobacco use cessation (Reid et al., 2016) or chronic pain (Chong and Bajwa, 2003), which complicates its use for reliably identifying depression (Gill et al., 2010). Lastly, the use of natural language processing (NLP) on clinical notes has great promise for classifying psychiatric disorders (Castro et al., 2015; Chen et al., 2018; Kho et al., 2011; Perlis et al., 2012; Zhou et al., 2015). However, the generalizability of these methods may be limited due to data sets that do not have the number or types of notes required or contain only deidentified structured data. Finally, most methods involve the exclusion of a sizeable number of patients with uncertain status, which prevents clinically relevant population-wide studies that include and classify all patients in the population.

This study examines a novel phenotyping algorithm for defining depression along a continuum using EHR data and evaluates their construct validity with other indicators of health that should correlate with depression. We focus on definitions based on ICD diagnosis codes and medication order data because these sources of data are more ubiquitously available than depression screening data and/or clinical notes and thus the resulting definitions are more widely generalizable.

## METHODS

### Study data and analysis

This study included de-identified Electronic Health Records (EHR) data obtained from January 1st, 2005 to September 30th, 2015 (10.75 years) for patients seen in the Geisinger Health System, an integrated health care system located in central Pennsylvania. The Geisinger system has a stable patient population whose EHRs have been collected in a central data warehouse and are available for clinical and research purposes (Borthwick et al., 2015; Boscarino et al., 2016; Carey et al., 2016; Maeng et al., 2017). The end date of the study period was chosen based on the transition from ICD-9 codes to ICD-10 codes in this hospital system. Adult patients 18 years or older at the beginning of the study, 90 years or younger at the end of the study, who had a Geisinger Primary Care Physician (PCP) at any point during the study period, and had at least one outpatient visit within the system during the study period were included in the study cohort (n=278,026) Demographic information, medication order histories, and details of all outpatient, Emergency Department (ED), and inpatient encounters were obtained for these patients. The study was approved by the Geisinger Internal Review Board as non-human subjects research.

We used domain knowledge of depression clinical care to implement an algorithm for partitioning all patients into one of five ordinal phenotype groups reflecting decreasing/increasing likelihood and/or severity of depression based on the available ICD-9 diagnosis codes and medication orders. To evaluate the convergent/divergent validity of these phenotype groups, we examined how they related to other health care related characteristics previously demonstrated to be associated with depression. These other characteristics include medical history, measures of health care utilization, depressive symptoms, and polygenic risk for depression. All analyses and visualizations described in further detail below were conducted in R (2017, R Core Team, Vienna, Austria) and GraphPad Prism 6 (La Jolla, CA).

### Depression phenotype algorithm

To implement the phenotype algorithm (Figure 1), patients were first grouped into “Major Depressive Disorder” (MDD) if they received a diagnosis of Major Depressive Disorder (ICD-9 codes: 296.20, 296.21, 296.22, 296.23, 296.24, 296.25, 296.26, 296.30, 296.31, 296.32, 296.33, 296.34, 296.35, 296.36, 296.82) one or more times as a discharge diagnosis during the study period. Second, the remaining patients who did not meet previous criteria were then grouped into “Other Depression” (OthDep) if they received a diagnosis of a depressive disorder not elsewhere classified (ICD9 code: 311) one or more times as a discharge diagnosis during the study period. Third, the remaining patients who did not meet any of these previous criteria were then grouped into “Antidepressants for Other Psychiatric Condition” (PsychRx) if they received one or more antidepressant medication orders (RxNorm classification “Antidepressant”) during the study period and received a concurrent discharge diagnosis for a psychiatric condition other than related to depression (IDC-9 codes: 290-319). Fourth, the remaining patients who did not meet any of these previous criteria but received one or more antidepressant medication orders were then grouped into “Antidepressants for Physical Condition” (NonPsychRx). Finally, all other remaining patients were grouped into “No Depression” (NoDep).

**Figure 1.**
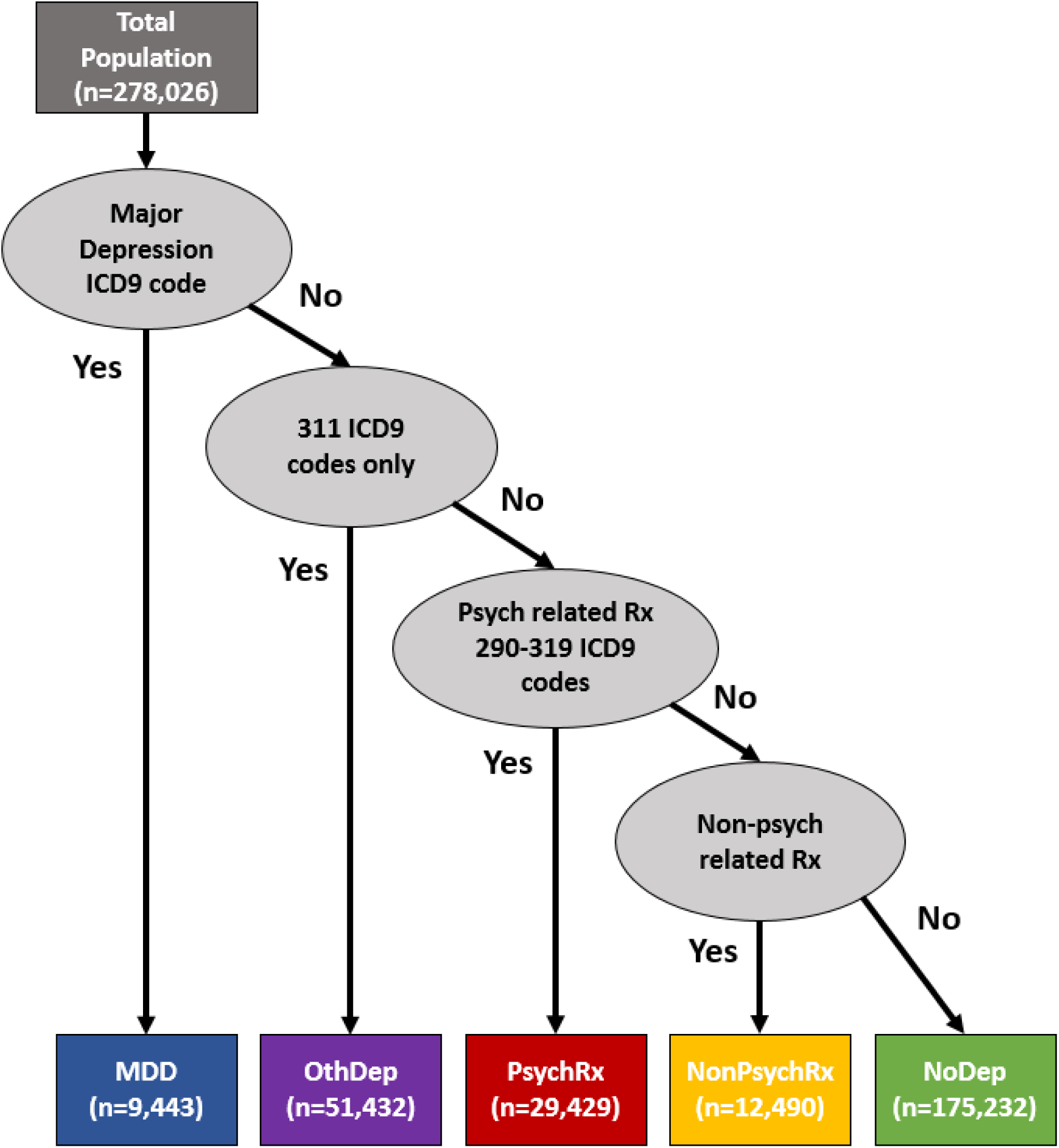
Methodological design with five mutually exclusive constructs including multiple depression phenotypes and one control group with no evidence of depression.

### Medical features

We evaluated several medical features that are known to be associated with depression, including medications, comorbidities and mortality. Depressed patients are often prescribed antipsychotics to augment their antidepressant medication (Wen et al., 2014), and antianxiety agents to treat anxiety, a common comorbid condition with depression (Verger et al., 2008). We determined the percent of patients in each group that had ever received at least one order for these medications as classified in RxNorm (“Antipsychotic” and “Antianxiety Agent”, respectively). Depressed patients are also known to have higher rates of suicidality (Hawton et al., 2013; Weitz et al., 2014) and mortality (Schulz et al., 2002; Wulsin et al., 1999) than non-depressed patients. We used the discharge diagnosis codes from encounter records to determine the percent of patients with ICD-9 codes for suicide and self-inflicted injury (E950 - E958). Substance abuse was captured using discharge diagnosis codes from encounter records to determine the percent of patients in each group that had ICD-9 codes for alcohol and drug abuse or dependence (303, 304, and 305). Because depressed patients have greater overall burden of medical comorbidities than non-depressed patients (Moussavi et al., 2007), we calculated the Charlson Comorbidity Index (CCI) score for each patient and compared the mean CCI across groups. The CCI contains 19 categories of serious comorbidities that predicts the 10-year mortality of patients (Deyo et al., 1992). Finally, we used the date of death to determine the percent mortality of each group during the study period.

### Healthcare utilization

Depression is associated with increased healthcare utilization (Bhattarai et al., 2013; Felleman et al., 2013; Possemato et al., 2013). As a measure of healthcare utilization, we calculated the average number of visits annually across different health care settings and compared these across the groups. To calculate the average number of visits in each healthcare setting per patient per year, we tallied the number of total visits in the outpatient, ED, non-psychiatric inpatient, and psychiatric inpatient settings during the entire study period for each group and divided this by the total number of patients in that group and the length of the study period (10.75 years), resulting in an estimated average yearly visit frequency.

### Depression symptoms

In 2012, Geisinger Health System began implementing universal screening for depression with the Patient Health Questionnaire 2 (PHQ-2). If patients endorse either of the two screener questions, they are then asked the additional 7 questions (PHQ-9). The PHQ-9 is a validated instrument for assessing current depression symptom severity and has a tiered rating scale based on total score (0: no depression, 1-9: mild, 10-14: moderate, 15-19: moderately severe, 20+: severe) (Kroenke et al., 2001b; Manea et al., 2015). For all patients that had one or more PHQ-2/9result in their EHR (n=170,618; 61.4%), we identified their maximum score and determined the percent of patients in each group with maximum scores in the different PHQ-2/9 scoring categories.

### Polygenic risk scores

Geisinger Health System has recruited over 90,000 patients to participate in a genetics study called MyCode (Carey et al., 2016). A subset of the study population described above have participated in this study, allowing us to calculate polygenic risk scores (PRS) for comparison across depression groups. Using publicly available summary results from a recently published genome-wide association study (GWAS) of MDD (Wray et al., 2018), we calculated PRS for each patient in our data set that had genetic data available (n=52,775; 19%). Polygenic risk scores for MDD were calculated using PRSice-2 (Euesden et al., 2015) at eight predetermined p-value thresholds: 1, 0.5, 0.1, 0.05, 0.01, 0.001, 0.0001, 0.000001. As results were similar across all p-value thresholds, we only report results for the nominal threshold of p=0.05.

## RESULTS

We developed and applied an algorithm for grouping patients based on ICD codes and medication orders into one of five ordinal groups, varying in likelihood and severity of depression, which we named from most to least likely/severe: “MDD”, “OthDep”, “PsychRx”, “NonPsychRx”, and “NoDep” (Figure 1). Of the total patient population, each group accounts for 3.4%, 18.5%, 10.6%, 4.5%, and 63.0%, respectively. Summary statistics on sex, race, marital status, age at beginning of the study and length between first and last encounters are shown for the total patient population and all five groups in Table 1. The sample is 54.0% female, 95.7% white, and 60.9% married, with a median age of 45 and median length between first and last encounter of just over 7 years. As expected, depression occurs more commonly in females, reflected by an increased Female to Male ratio in each “affected” group when compared with the NoDep group. Patients in the MDD and OthDep groups are less likely to be “Married” compared with those in the NoDep, PsychRx, and NonPsychRx groups. Age at the beginning of the study does not differ substantially between most groups (median: 45, third quartile: 57 years), although it is slightly older for those in the NonPsychRx group (median: 48, third quartile: 60 years). The length between first and last encounter varied across groups, with a much higher percentage of patients in one of the four “affected” groups having at least 7 years of observation than those in the NoDep group. It is also of note that the vast majority of patients in the MDD and OthDep groups received at least one antidepressant medication order, 94.0% and 89.8%, respectively.

**Table 1.** Characteristics of the total sample and five depression phenotype groups

|  |  | **Total**<br>(n=278,026) |  | **MDD**<br>(n=9,443) |  | **OthDep**<br>(n=51,432) |  | **PsychRx**<br>(n=29,429) |  | **NonPsychRx**<br>(n=12,490) |  | **NoDep**<br>(n=175,232) |  |
| --- | --- | --- | --- | --- | --- | --- | --- | --- | --- | --- | --- | --- | --- |
|  |  | n | % | n | % | n | % | n | % | n | % | n | % |
| **Sex** | Male | 127919 | 46.0 | 2853 | 30.2 | 16580 | 32.2 | 11356 | 38.6 | 4564 | 36.5 | 92566 | 52.8 |
|  | Female | 150107 | 54.0 | 6590 | 69.8 | 34852 | 67.8 | 18073 | 61.4 | 7926 | 63.5 | 82666 | 47.2 |
| **Race** | White | 266043 | 95.7 | 9264 | 98.1 | 50053 | 97.3 | 28818 | 97.9 | 12098 | 96.9 | 165810 | 94.6 |
|  | Black | 7168 | 2.6 | 124 | 1.3 | 990 | 1.9 | 402 | 1.4 | 244 | 2.0 | 5408 | 3.1 |
|  | Asian | 2281 | 0.8 | 22 | 0.2 | 133 | 0.3 | 85 | 0.3 | 71 | 0.6 | 1970 | 1.1 |
|  | Native | 1384 | 0.5 | 23 | 0.2 | 153 | 0.3 | 84 | 0.3 | 44 | 0.4 | 1080 | 0.6 |
|  | Other | 1150 | 0.4 | 10 | 0.1 | 103 | 0.2 | 40 | 0.1 | 33 | 0.3 | 964 | 0.6 |
| **Marital Status** | Married | 169252 | 60.9 | 4827 | 51.1 | 27362 | 53.2 | 17606 | 59.8 | 7997 | 64.0 | 111467 | 63.6 |
|  | Single | 57848 | 20.8 | 1924 | 20.4 | 10297 | 20.0 | 6008 | 20.4 | 1932 | 15.5 | 37687 | 21.5 |
|  | Widowed | 21002 | 7.6 | 831 | 8.8 | 5268 | 10.2 | 2161 | 7.3 | 1183 | 9.5 | 11559 | 6.6 |
|  | Divorced | 24480 | 8.8 | 1480 | 15.7 | 6809 | 13.2 | 2997 | 10.2 | 1159 | 9.3 | 12035 | 6.9 |
|  | Other | 5437 | 2.0 | 381 | 4.0 | 1696 | 3.3 | 657 | 2.2 | 219 | 1.8 | 2484 | 1.4 |
| **Age at beginning of study** | 18-30 | 60093 | 21.6 | 1976 | 20.9 | 10680 | 20.8 | 7158 | 24.3 | 1987 | 15.9 | 38292 | 21.9 |
|  | 31-45 | 85036 | 30.6 | 3152 | 33.4 | 16237 | 31.6 | 10096 | 34.3 | 3498 | 28.0 | 52053 | 29.7 |
|  | 46-65 | 98727 | 35.5 | 3448 | 36.5 | 18424 | 35.8 | 9343 | 31.8 | 4804 | 38.5 | 62708 | 35.8 |
|  | 66+ | 34170 | 12.3 | 867 | 9.2 | 6091 | 11.8 | 2832 | 9.6 | 2201 | 17.6 | 22179 | 12.6 |
| **Length between first and last encounters** | < 1 yr | 36523 | 13.1 | 402 | 4.3 | 2257 | 4.4 | 1018 | 3.5 | 503 | 4.0 | 32343 | 18.5 |
|  | 1-4 yrs | 60817 | 21.9 | 1221 | 12.9 | 9370 | 18.2 | 5081 | 17.3 | 2157 | 17.3 | 42988 | 24.5 |
|  | 4-7 yrs | 41172 | 14.8 | 1243 | 13.2 | 7908 | 15.4 | 4583 | 15.5 | 1846 | 14.8 | 25592 | 14.6 |
|  | 7-10.75 yrs | 139514 | 50.2 | 6577 | 69.6 | 31897 | 62.0 | 18747 | 63.7 | 7984 | 63.9 | 74309 | 42.4 |

### Medication orders and comorbidities

The percent of patients that received antipsychotic (Figure 2A) or antianxiety agents (Figure 2B) decreased monotonically across the five groups from a high for MDD to a low for NoDep. Approximately 44% of MDD compared to 5.6% of NoDep patients were prescribed antipsychotic medications, while 30% of MDD compared to 6% of NoDep patients were prescribed antianxiety medications. A similar monotonic decreasing pattern was observed across the five phenotype groups for the percent of patients diagnosed with substance use and abuse (Figure 2C), although here the decline was more step-wise. The decreasing percentage across groups was more dramatic for suicide related diagnosis codes (Figure 2D). Almost all suicide related diagnosis codes were observed in MDD and OthDep patients, while the percent of patient with such codes was well below 1% in the other three groups. Finally, the mean CCI scores (Figure 2E) and overall mortality (Figure 2F) generally decreased across the five groups as expected with the marked exception of the NonPsychRx group. Interestingly, the highest mean CCI and mortality rates were observed for patients in the NonPsychRx group.

**Figure 2.**
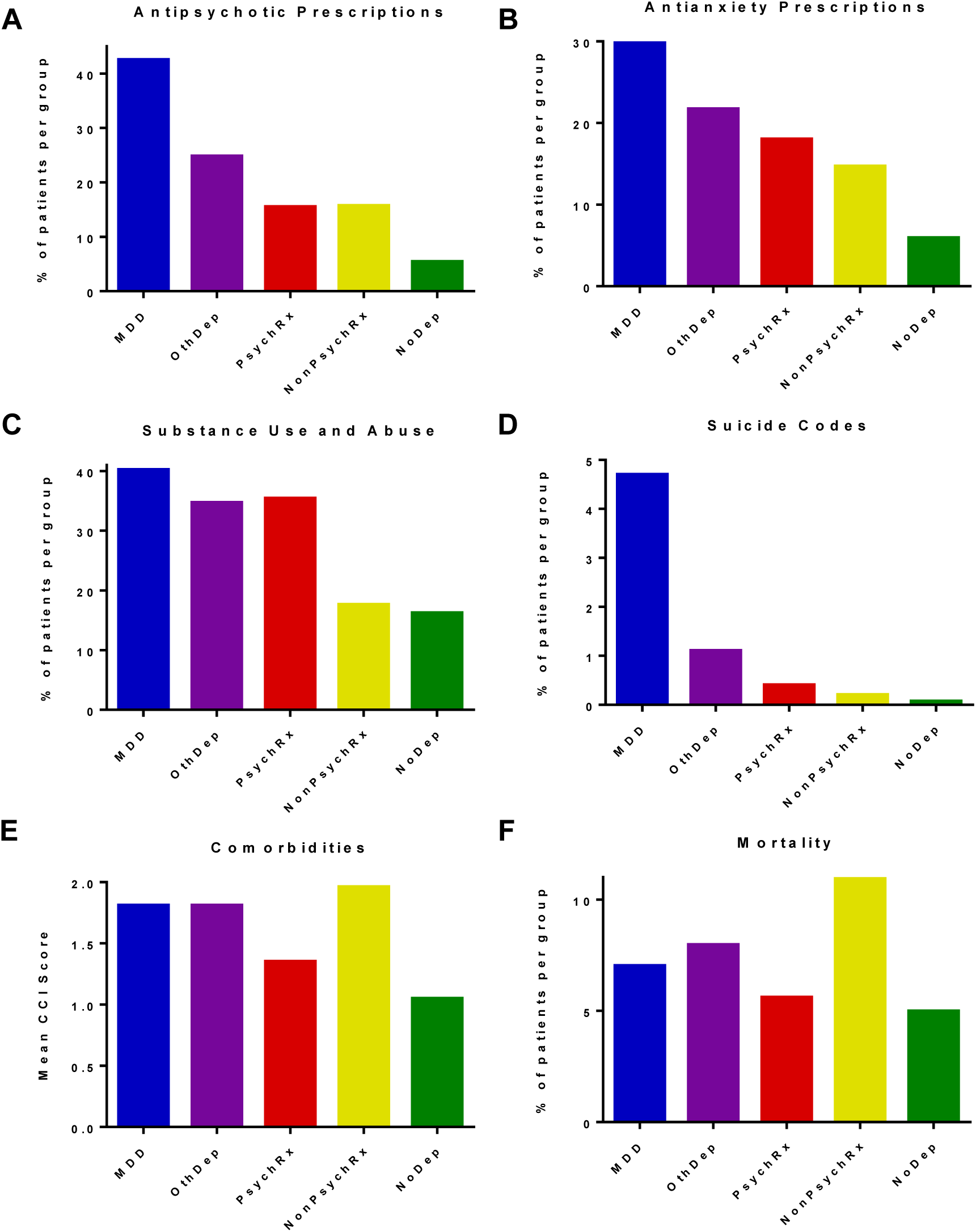
Medical features across the five depression phenotype groups. (A) Antipsychotic and (B) antianxiety medication prescriptions, (C) substance use disorder or dependence codes, (D) suicide codes, (E) mean Charlson Comorbidity Index scores, and (F) mortality.

### Healthcare utilization

The most striking difference in utilization across the five phenotype groups was observed for psychiatric inpatient visits (Figure 3A). MDD patients had an average of 0.03 such visits/year, which was 10 times greater than for OthDep patients, while the rates were negligible for the other three groups. The average number of visits/year decreased nearly monotonically across the five phenotype groups from MDD to NoDep for nonpsychiatric inpatient visits (ranging from 0.13 to 0.03 visits/year, respectively), ED visits (ranging from 0.24 to 0.06 visits/year, respectively), and outpatient visits (ranging from 5.4 to 2.0 visits/year, respectively) (Figures 3B, 3C, and 3D). A marked exception to this trend was observed for the NonPsychRx group which had higher average visits/year for all three visit types than both the PsychRx and NoDep groups.

**Figure 3.**
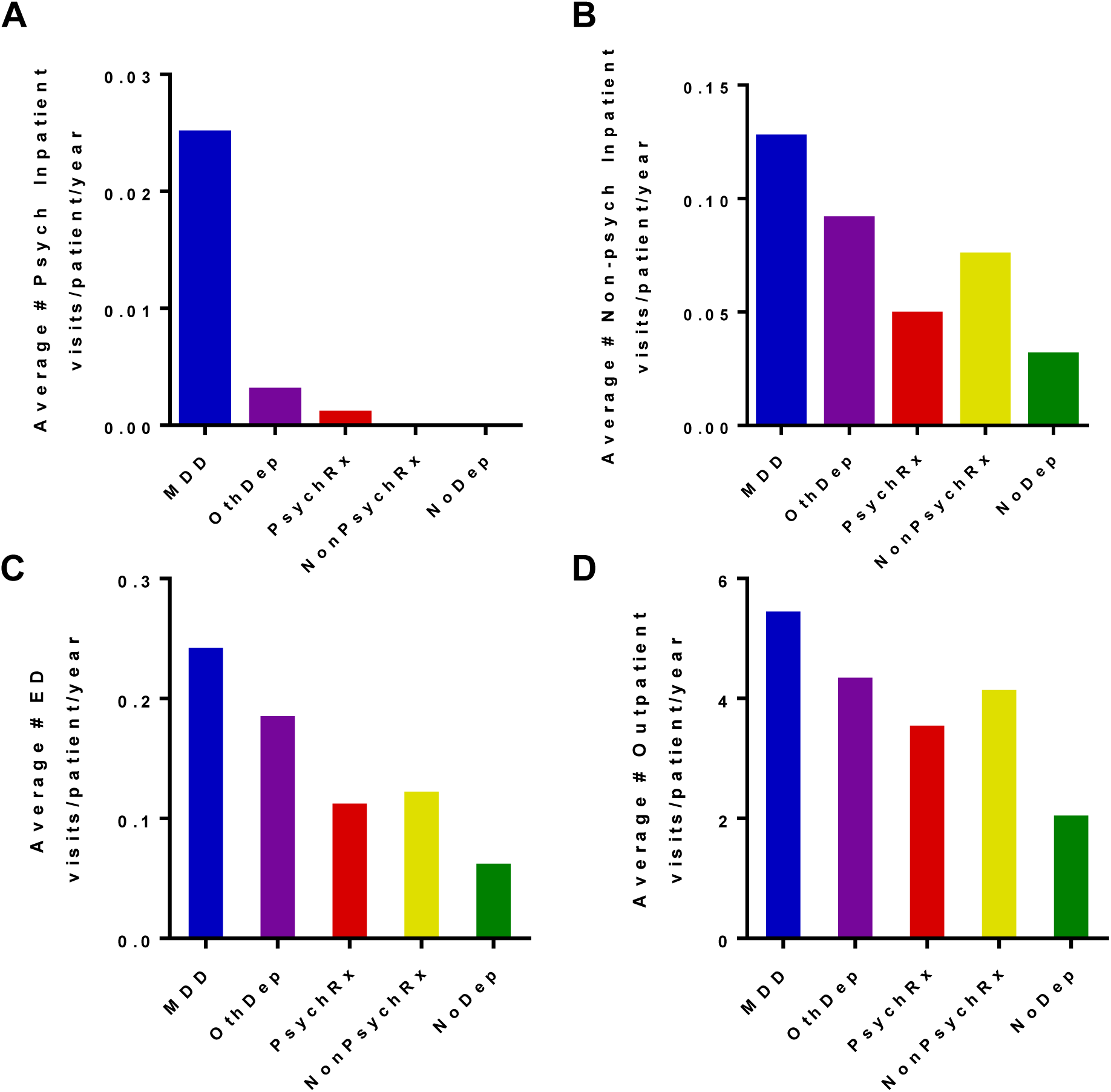
Healthcare utilization patterns for the five depression phenotype groups. (A) Psychiatric inpatient visit frequency; (B) Non-psychiatric inpatient visit frequency; (C) Emergency Department (ED) visit frequency; (D) Outpatient visit frequency.

### Depression symptoms

A subset of the study population was screened at least once with the Patient Health Questionnaire (PHQ), a well validated depression measure (Figure 4). Patients in the MDD group were the most likely to have been screened (72.7%, n=6865), followed by those in the OthDep (70.2%; n=36,091), PsychRx (67.6%; n=19,898), NonPsychRx (65.2%; n=8141), and finally NoDep (56.9%; n=99,623). Again, we observed a monotonic decrease across the five phenotype groups in the percentage of patients with maximum PHQ scores in the higher categories indicating mild, moderate, moderately severe and severe depression. This was accompanied by a complimentary monotonic decrease in the percentage of patients with a maximum score of 0 indicating no depression. Nearly a third of the MDD group had a maximum PHQ score of 10 or higher, while approximately only 30% had a maximum score of 0. Conversely, almost 80% of patients in the NoDep group had a maximum score of 0, while only 2.9% scored 10 or higher.

**Figure 4.**
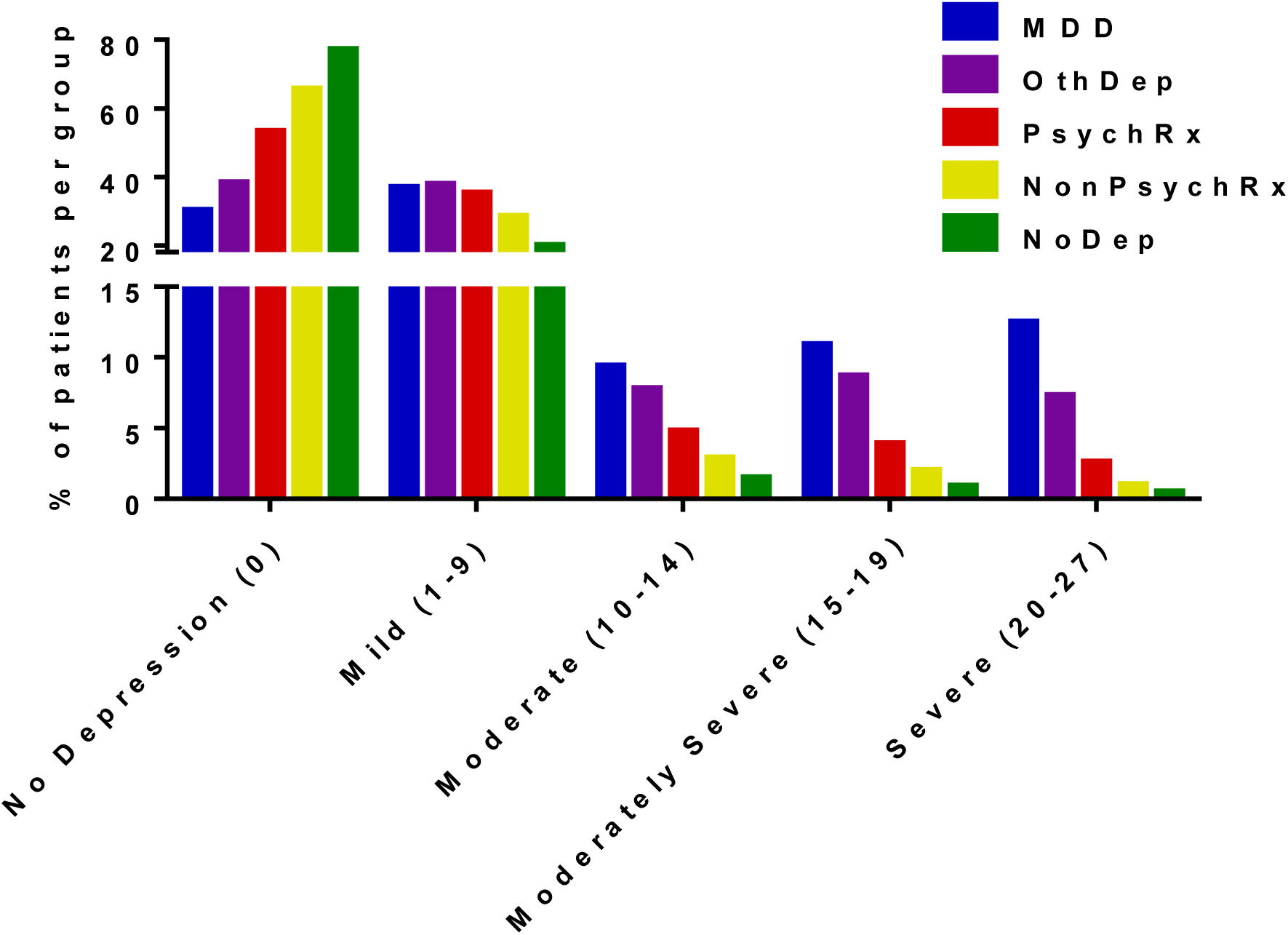
Depression symptoms across the five depression phenotype groups. Percentage of patients per group whose maximum score on the Patient Health Questionnaire 2 or 9 (PHQ-9/2) fell in the categories shown.

### Polygenic risk scores

As depression has a prominent heritable component (Hamet and Tremblay, 2005), we used polygenic risk scores (PRS) derived from an external GWAS (Wray et al., 2018) to further validate our phenotype groups for those patients in this study who had available genome-wide genotype data. As shown in Figure 5, we found a gradient of increased PRSs across the five phenotype groups, with the highest PRSs seen in the MDD group and the greatest difference seen between the MDD and NoDep groups (P < 2.2×10^-16^, R2 =0.8%).

**Figure 5.**
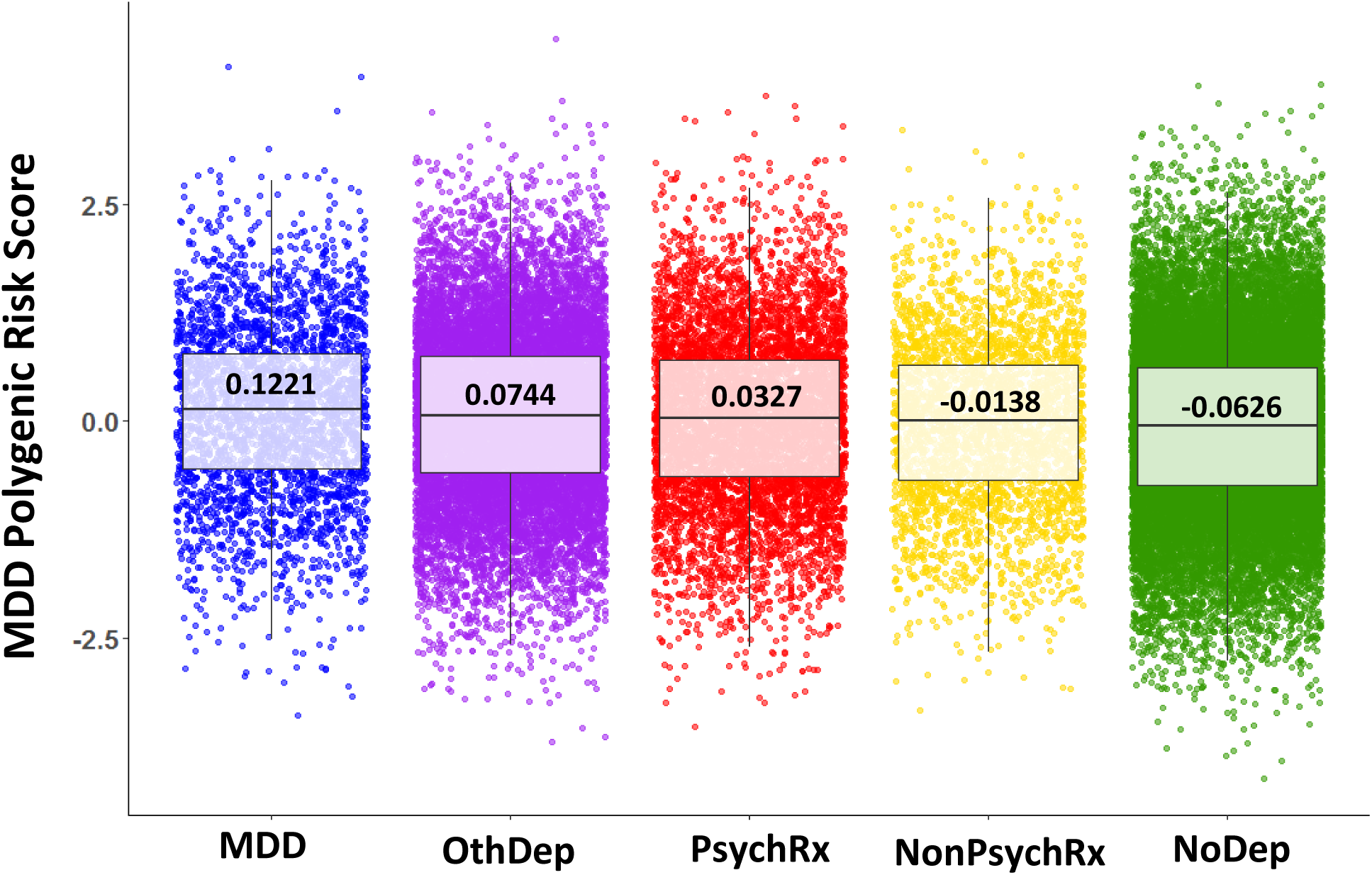
Scaled Polygenic Risk Scores for Major Depressive Disorder correlate with and distinguish between depression constructs when compared with NoDep group. P-value threshold 0.05, includes 20,962 SNPs. Comparing NoDep to both MDD groups combined results in R2=0.008. Welch two sample t-test between each group and NoDep results in statistically significantly differences in each comparison. Median values displayed in bold at center of boxplots overlaid on jittered dot plots.

## DISCUSSION

We present a novel electronic phenotype algorithm using ICD codes and medication orders in EHR data, resulting in five mutually exclusive and ordinal groups for categorizing patients with depression across an entire population. These phenotype groups demonstrate convergent validity as assessed by significant differences between them in a range of clinically relevant characteristics that are expected to correlate with depression (Kamiya et al., 2013; Yan et al., 2011), including medical features (such as treatments (Kessing et al., 2017), comorbidities (Gallo et al., 2016), and mortality), healthcare utilization (Bhattarai et al., 2013), depression screening results, and polygenic risk scores. Interestingly, these clinically relevant characteristics tended to vary in a “dose-response” like fashion across the phenotype groups, consistent with the fact that the groups defined by increasingly more stringent criteria identified patients with increasing likelihood and/or severity of depression. The one notable exception to this pattern was observed for patients in the NonPsychRx group who tended to have higher rates of comorbidity, non-psychiatric related hospital encounters and overall mortality than might be expected, which may have been due to the fact that patients in this group were older on average than those in the other groups. Overall, these findings demonstrate that it is possible to use diagnostic codes and medications orders in EHR data to validly categorize patients with respect to depression likelihood/severity across the entire population.

With the wide spread adoption of EHRs, there is growing interest in identifying patients with depression for subsequent research using data available from the EHRs. As a result, a number of previous studies have attempted to do this employing a variety of methods that use, in isolation or in combination, ICD9 diagnosis codes, antidepressant prescription orders, and NLP of progress notes. Many of these studies have not validated their algorithms (Anderson et al., 2015; Gill et al., 2008; Kobus et al., 2013; Madden et al., 2016; Mayer et al., 2017). There are a few studies, however, that have attempted to validate their algorithms by comparing identified cases and controls against a putative gold-standard diagnosis established by expert review of a relatively limited number of medical charts per group (Castro et al., 2015; Huang et al., 2014; Perlis et al., 2012). Our approach differs from these previous efforts in that we sought to establish convergent rather than criterion validation of our algorithm, and we sought to characterize and compare the full range of depression in the entire sample rather than a small sub-sample of putative cases and controls which presumably represent the extremes of the phenotype in the population. In addition, other than one study which used genetic data to try and validate an algorithm for defining a bipolar disorder phenotype (Chen et al., 2018), our study is the only one as far as we know to use genetic data to help validate an algorithm for defining depression.

This study has several limitations. First, with regard to our examination of healthcare utilization patterns and medical features, we made the assumption that patients contributed follow-up over the entire study period. We did this to simplify our estimates, though it is possible that a number of patients entered or left the Geisinger service area during the study period. Second, with regard to our algorithm, we categorized patients based on the most severe criteria met over the study period and assumed they belonged to that group for the entire period. This lifetime approach does not allow for change over time and does not take full advantage of this rich longitudinal data, which may be interesting to study using longitudinal data analysis methods in the future. Our algorithm was also based on ICD-9 codes, as opposed to ICD-10 codes which are the current version in use for diagnosis and healthcare billing. Fortunately, there are many resources available for mapping between ICD-9 and ICD-10 codes that can be leveraged to create an equivalent list of ICD-10 codes to the list and codes used here (CMS, 2015; ICD10Data.com, n.d.). Third, the sample under study consisted of predominantly white patients from rural communities in central Pennsylvania. As a result, there may be questions about the generalizability of our approach to other more diverse and urban populations. In addition, there are other healthcare systems in the service area and thus the data we used to categorize patients may not have captured all medical utilization. Claims data may provide an additional source of information about healthcare utilization that would be useful in future studies.

This study also benefits from several significant strengths. One of the major strengths of this study is the large longitudinal data from a stable patient population in an integrated health care system. With extended and extensive data available from multiple health care settings (primary, emergency, and hospital care), we were able to convergently validate our simple phenotype algorithm by comparison with a range of different measures. Additionally, our algorithm allows for classification of all patients in a given population across a spectrum of depression severity, as opposed to other approaches which often seek to classify only the extremes of the population to the exclusion of patients with uncertain binary case versus control status (Castro et al., 2015). Finally, our algorithm uses only diagnosis codes and medication data to validly classify patients with regard to depression and therefore may be more generalizable to other systems that do not have ready access to data on screening measures or from natural language processing of clinician notes.

The electronic phenotype algorithm presented here provides a simple and valid model for defining patients with varying likelihood or severities of depression and may be useful for researchers who wish to examine the effects of depression in entire patient populations. The five mutually exclusive, ordinal groups demonstrate differences that are expected to correlate with depression. We found correlations with treatments, comorbidities, mortality, utilization patterns, depression screening instruments for symptom severity, and polygenic risk scores. These constructs are generalizable to any data set that has both diagnosis codes and medication orders. Ultimately, the definition of depression phenotypes will depend upon the goal of the research, and we present one possible method that demonstrates convergent validity, generalizability, and inclusivity.

## ACKNOWLEDGEMENTS, COMPETING INTERESTS, FUNDING

### Funding

This work was supported by the National Institutes of Health (T32MH014592-41 Psychiatric Epidemiology Training Program)

## REFERENCES

Affolter, K., Gligorich, K., Samadder, N.J., Samowitz, W.S., Curtin, K., 2017. Feasibility of Large-Scale Identification of Sessile Serrated Polyp Patients Using Electronic Records: A Utah Study. Dig. Dis. Sci. 6, 1455–1463. https://doi.org/10.1007/s10620-017-4543-9

Anderson, H.D., Pace, W.D., Brandt, E., Nielsen, R.D., Allen, R.R., Libby, A.M., West, D.R., Valuck, R.J., 2015. Monitoring suicidal patients in primary care using electronic health records. J. Am. Board Fam. Med. 28, 65–71. https://doi.org/10.3122/jabfm.2015.01.140181

Ang, D.C., Choi, H., Kroenke, K., Wolfe, F., 2005. Comorbid depression is an independent risk factor for mortality in patients with rheumatoid arthritis. J. Rheumatol. 32, 1013–1019.

Bhattarai, N., Charlton, J., Rudisill, C., Gulliford, M.C., 2013. Prevalence of depression and utilization of health care in single and multiple morbidity: a population-based cohort study. Psychol. Med. 43, 1423–31. https://doi.org/10.1017/S0033291712002498

Borthwick, K., Smelser, D., Bock, J., Elmore, J., Ryer, E., Ye, Z., Pacheco, J., Carrell, D., Michalkiewicz, M., Thompson, W., Pathak, J., Bielinski, S., Denny, J., Linneman, J., Peissig, P., Kho, A., Gottesman, O., Parmar, H., Kullo, I., McCarty, C., Böttinger, E., Larson, E., Jarvik, G., Harley, J., Bajwa, T., Franklin, D., Carey, D., Kuivaniemi, H., Tromp, G., 2015. Ephenotyping for Abdominal Aortic Aneurysm in the Electronic Medical Records and Genomics (emerge) Network: Algorithm Development and Konstanz Information Miner Workflow. Int. J. Biomed. Data Min. 4, 1–8. https://doi.org/10.4172/2090-4924.1000113

Boscarino, J.A., Kirchner, H.L., Pitcavage, J.M., Nadipelli, V.R., Ronquest, N.A., Fitzpatrick, M.H., Han, J.J., 2016. Factors associated with opioid overdose: a 10-year retrospective study of patients in a large integrated health care system. Subst. Abuse Rehabil. 7, 131–141. https://doi.org/10.2147/SAR.S108302

Carey, D.J., Fetterolf, S.N., Davis, F.D., Faucett, W.A., Kirchner, H.L., Mirshahi, U., Murray, M.F., Smelser, D.T., Gerhard, G.S., Ledbetter, D.H., 2016. The Geisinger MyCode community health initiative: An electronic health record-linked biobank for precision medicine research. Genet. Med. 18, 906–913. https://doi.org/10.1038/gim.2015.187

Castro, V.M., Minnier, J., Murphy, S.N., Kohane, I., Churchill, S.E., Gainer, V., Cai, T., Hoffnagle, A.G., Dai, Y., Block, S., Weill, S.R., Nadal-Vicens, M., Pollastri, A.R., Rosenquist, J.N., Goryachev, S., Ongur, D., Sklar, P., Perlis, R.H., Smoller, J.W., Lee, P.H., Stahl, E.A., Purcell, S.M., Ruderfer, D.M., Charney, A.W., Roussos, P., Pato, C., Pato, M., Medeiros, H., Sobel, J., Craddock, N., Jones, I., Forty, L., DiFlorio, A., Green, E., Jones, L., Dunjewski, K., Landén, M., Hultman, C., Juréus, A., Bergen, S., Svantesson, O., McCarroll, S., Moran, J., Chambert, K., Belliveau, R.A., 2015. Validation of electronic health record phenotyping of bipolar disorder cases and controls. Am. J. Psychiatry 172, 363–372. https://doi.org/10.1176/appi.ajp.2014.14030423

Celano, C.M., Huffman, J.C., 2011. Depression and Cardiac Disease. Cardiol. Rev. 19, 130–142. https://doi.org/10.1097/CRD.0b013e31820e8106

Chen, C.-Y., Lee, P.H., Castro, V.M., Minnier, J., Charney, A.W., Stahl, E.A., Ruderfer, D.M., Murphy, S.N., Gainer, V., Cai, T., Jones, I., Pato, C.N., Pato, M.T., Landén, M., Sklar, P., Perlis, R.H., Smoller, J.W., 2018. Genetic validation of bipolar disorder identified by automated phenotyping using electronic health records. Transl. Psychiatry 8, 86. https://doi.org/10.1038/s41398-018-0133-7

Chong, M.S., Bajwa, Z.H., 2003. Diagnosis and treatment of neuropathic pain. J. Pain Symptom Manage. 25, S4–S11.

CMS, 2015. CMS Chronic Conditions Data Warehouse (CCW) CCW Condition Algorithms. Chronic Cond. Algorithms 72, 2015. https://doi.org/10.1002/pds.2329/pdf

Cole, A.M., Stephens, K.A., Keppel, G.A., Estiri, H., Baldwin, L.-M., 2016. Extracting Electronic Health Record Data in a Practice-Based Research Network: Lessons Learned from Collaborations with Translational Researchers. eGEMs (Generating Evid. Methods to Improv. patient outcomes) 4, 4. https://doi.org/10.13063/2327-9214.1206

Conway, M., Berg, R.L., Carrell, D., Denny, J.C., Kho, A.N., Kullo, I.J., Linneman, J.G., Pacheco, J.A., Peissig, P., Rasmussen, L., Weston, N., Chute, C.G., Pathak, J., 2011. Analyzing the heterogeneity and complexity of Electronic Health Record oriented phenotyping algorithms. AMIA Annu Symp Proc 2011, 274–283. https://doi.org/PMC3243189

Deyo, R.A., Cherkin, D.C., Ciol, M.A., 1992. Adapting a clinical comorbidity index for use with ICD-9-CM administrative databases. J. Clin. Epidemiol. 45, 613–619. https://doi.org/10.1016/0895-4356(92)90133-8

Dockery, L., Jeffery, D., Schauman, O., Williams, P., Farrelly, S., Bonnington, O., Gabbidon, J., Lassman, F., Szmukler, G., Thornicroft, G., Clement, S., 2015. Stigma- and non-stigma-related treatment barriers to mental healthcare reported by service users and caregivers. Psychiatry Res. 228, 612–619. https://doi.org/10.1016/j.psychres.2015.05.044

Euesden, J., Lewis, C.M., O’Reilly, P.F., 2015. PRSice: Polygenic Risk Score software. Bioinformatics 31, 1466–1468. https://doi.org/10.1093/bioinformatics/btu848

Felleman, B.I., Athenour, D.R., Ta, M.T., Stewart, D.G., 2013. Behavioral health services influence medical treatment utilization among primary care patients with comorbid substance use and depression. J. Clin. Psychol. Med. Settings 20, 415–26. https://doi.org/10.1007/s10880-013-9367-y

Fiest, K.M., Jette, N., Quan, H., St Germaine-Smith, C., Metcalfe, A., Patten, S.B., Beck, C.A., 2014. Systematic review and assessment of validated case definitions for depression in administrative data. BMC Psychiatry 14, 1–11. https://doi.org/10.1186/s12888-014-0289-5

Gallo, J.J., Hwang, S., Joo, J.H., Bogner, H.R., Morales, K.H., Bruce, M.L., Reynolds, C.F., 2016. Multimorbidity, Depression, and Mortality in Primary Care: Randomized Clinical Trial of an Evidence-Based Depression Care Management Program on Mortality Risk. J. Gen. Intern. Med. 31, 380–386. https://doi.org/10.1007/s11606-015-3524-y

Gill, J.M., Chen, Y.X., Lieberman, M.I., 2008. Management of depression in ambulatory care for patients with medical co-morbidities: a study from a national Electronic Health Record (EHR) network. Int. J. Psychiatry Med. 38, 203–15. https://doi.org/10.2190/PM.38.2.g

Gill, J.M., Klinkman, M.S., Chen, Y.X., 2010. Antidepressant medication use for primary care patients with and without medical comorbidities: a national electronic health record (EHR) network study. J. Am. Board Fam. Med. JABFM 23, 499–508. https://doi.org/10.3122/jabfm.2010.04.090299

Green, A.C., Hunt, C., Stain, H.J., 2012. The delay between symptom onset and seeking professional treatment for anxiety and depressive disorders in a rural Australian sample. Soc. Psychiatry Psychiatr. Epidemiol. 47, 1475–1487. https://doi.org/10.1007/s00127-011-0453-x

Hamet, P., Tremblay, J., 2005. Genetics and genomics of depression. Metabolism. 54, 10–15. https://doi.org/10.1016/j.metabol.2005.01.006

Hawton, K., Casañasi Comabella, C., Haw, C., Saunders, K., 2013. Risk factors for suicide in individuals with depression: A systematic review. J. Affect. Disord. 147, 17–28. https://doi.org/10.1016/j.jad.2013.01.004

Huang, S.H., LePendu, P., Iyer, S. V, Tai-Seale, M., Carrell, D., Shah, N.H., 2014. Toward personalizing treatment for depression: predicting diagnosis and severity. J. Am. Med. Inform. Assoc. 21, 1–7. https://doi.org/10.1136/amiajnl-2014-002733

ICD10Data.com, n.d. Convert ICD-9-CM Codes to ICD-10-CM codes [WWW Document]. URL https://www.icd10data.com/Convert

Jin, H., Wu, S., Vidyanti, I., Di Capua, P., Wu, B., 2015. Predicting Depression among Patients with Diabetes Using Longitudinal Data. A Multilevel Regression Model. Methods Inf Med 54, 553–559. https://doi.org/10.3414/ME14-02-0009

Kalia, M., 2005. Neurobiological basis of depression: An update. Metabolism. 54, 24–27. https://doi.org/10.1016/j.metabol.2005.01.009

Kamiya, Y., Doyle, M., Henretta, J.C., Timonen, V., 2013. Depressive symptoms among older adults: the impact of early and later life circumstances and marital status. Aging Ment. Health 17, 349–57. https://doi.org/10.1080/13607863.2012.747078

Kessing, L.V., Willer, I., Andersen, P.K., Bukh, J.D., 2017. Rate and predictors of conversion from unipolar to bipolar disorder: A systematic review and meta-analysis. Bipolar Disord. 19, 324–335. https://doi.org/10.1111/bdi.12513

Kho, A.N., Pacheco, J.A., Peissig, P.L., Rasmussen, L., Newton, K.M., Weston, N., Crane, P.K., Pathak, J., Chute, C.G., Bielinski, S.J., Kullo, I.J., Li, R., Manolio, T.A., Chisholm, R.L., Denny, J.C., 2011. Electronic Medical Records for Genetic Research: Results of the eMERGE Consortium. Sci. Transl. Med. 3, 79re1–79re1. https://doi.org/10.1126/scitranslmed.3001807

Kobus, A.M., Harman, J.S., Do, H.D., Garvin, R.D., 2013. Challenges to depression care documentation in an EHR. Fam. Med. 45, 268–271.

Kroenke, K., Spitzer, R.L., Williams, J.B.W., 2001a. The PHQ-9. J. Gen. Intern. Med. 16, 606–613. https://doi.org/10.1046/j.1525-1497.2001.016009606.x

Kroenke, K., Spitzer, R.L., Williams, J.B.W., 2001b. The PHQ-9: Validity of a brief depression severity measure. J. Gen. Intern. Med. 16, 606–613. https://doi.org/10.1046/j.1525-1497.2001.016009606.x

Madden, J.M., Lakoma, M.D., Rusinak, D., Lu, C.Y., Soumerai, S.B., 2016. Missing clinical and behavioral health data in a large electronic health record (EHR) system. J. Am. Med. Informatics Assoc. 23, 1143–1149. https://doi.org/10.1093/jamia/ocw021

Maeng, D.D., Han, J.J., Fitzpatrick, M.H., Boscarino, J.A., 2017. Patterns of health care utilization and cost before and after opioid overdose: findings from 10-year longitudinal health plan claims data. Subst. Abuse Rehabil. 8, 57–67. https://doi.org/10.2147/SAR.S135884

Manea, L., Gilbody, S., McMillan, D., 2015. A diagnostic meta-analysis of the Patient Health Questionnaire-9 (PHQ-9) algorithm scoring method as a screen for depression. Gen. Hosp. Psychiatry 37, 67–75. https://doi.org/10.1016/j.genhosppsych.2014.09.009

Maurer, D.M., 2012. Screening for Depression. Am. Fam. Physician 85, 139–144.

Mayer, M.A., Gutierrez-Sacristan, A., Leis, A., De La Peña, S., Sanz, F., Furlong, L.I., 2017. Using Electronic Health Records to Assess Depression and Cancer Comorbidities. Stud. Health Technol. Inform. 235, 236–240. https://doi.org/10.3233/978-1-61499-753-5-236

McDonald, H.I., Shaw, C., Thomas, S.L., Mansfield, K.E., Tomlinson, L.A., Nitsch, D., 2016. Methodological challenges when carrying out research on CKD and AKI using routine electronic health records. Kidney Int. 90, 943–949. https://doi.org/10.1016/j.kint.2016.04.010

Morley, K.I., Wallace, J., Denaxas, S.C., Hunter, R.J., Patel, R.S., Perel, P., Shah, A.D., Timmis, A.D., Schilling, R.J., Hemingway, H., 2014. Defining Disease Phenotypes Using National Linked Electronic Health Records: A Case Study of Atrial Fibrillation. PLoS One 9. https://doi.org/10.1371/journal.pone.0110900

Moussavi, S., Chatterji, S., Verdes, E., Tandon, A., Patel, V., Ustun, B., 2007. Depression, chronic diseases, and decrements in health: results from the World Health Surveys. Lancet 370, 851–858. https://doi.org/10.1016/S0140-6736(07)61415-9

Nadkarni, G.N., Gottesman, O., Linneman, J.G., Chase, H., Berg, R.L., Farouk, S., Nadukuru, R., Lotay, V., Ellis, S., Hripcsak, G., Peissig, P., Weng, C., Bottinger, E.P., 2014. Development and validation of an electronic phenotyping algorithm for chronic kidney disease. AMIA Annu. Symp. Proc. 2014, 907–16.

Newton-Dame, R., McVeigh, K.H., Schreibstein, L., Perlman, S., Lurie-Moroni, E., Jacobson, L., Greene, C., Snell, E., Thorpe, L.E., 2016. Design of the New York City Macroscope: Innovations in Population Health Surveillance Using Electronic Health Records. eGEMs 4. https://doi.org/10.13063/2327-9214.1265

Newton, K.M., Peissig, P.L., Kho, A.N., Bielinski, S.J., Berg, R.L., Choudhary, V., Basford, M., Chute, C.G., Kullo, I.J., Li, R., Pacheco, J.A., Rasmussen, L. V., Spangler, L., Denny, J.C., 2013. Validation of electronic medical record-based phenotyping algorithms: Results and lessons learned from the eMERGE network. J. Am. Med. Informatics Assoc. 20, 147–154. https://doi.org/10.1136/amiajnl-2012-000896

Pathak, J., Simon, G., Li, D., Biernacka, J.M., Jenkins, G.J., Chute, C.G., Hall-Flavin, D.K., Weinshilboum, R.M., 2014. Detecting Associations between Major Depressive Disorder Treatment and Essential Hypertension using Electronic Health Records. AMIA Jt. Summits Transl. Sci. proceedings. AMIA Jt. Summits Transl. Sci. 2014, 91–96.

Perlis, R.H., Iosifescu, D. V., Castro, V.M., Murphy, S.N., Gainer, V.S., Minnier, J., Cai, T., Goryachev, S., Zeng, Q., Gallagher, P.J., Fava, M., Weilburg, J.B., Churchill, S.E., Kohane, I.S., Smoller, J.W., 2012. Using electronic medical records to enable large-scale studies in psychiatry: treatment resistant depression as a model. Psychol. Med. 42, 41–50. https://doi.org/10.1017/S0033291711000997

Possemato, K., Bishop, T.M., Willis, M.A., Lantinga, L.J., 2013. Healthcare utilization and symptom variation among veterans using behavioral telehealth center services. J. Behav. Heal. Serv. Res. 40, 416–426. https://doi.org/10.1007/s11414-013-9338-y

Pouwer, F., Nefs, G., Nouwen, A., 2013. Adverse Effects of Depression on Glycemic Control and Health Outcomes in People with Diabetes. Endocrinol. Metab. Clin. North Am., Endocrine and Neuropsychiatric Disorders 42, 529–544. https://doi.org/10.1016/j.ecl.2013.05.002

Reid, R.D., Pritchard, G., Walker, K., Aitken, D., Mullen, K.-A., Pipe, A.L., 2016. Managing smoking cessation. Can. Med. Assoc. J. 188, E484–E492. https://doi.org/10.1503/cmaj.151510

Sallakh, M.A. Al, Vasileiou, E., Rodgers, S.E., Lyons, R.A., Sheikh, A., Davies, G.A., 2017. Defining asthma and assessing asthma outcomes using electronic health record data: a systematic scoping review. Eur. Respir. J. 49, 1700204. https://doi.org/10.1183/13993003.00204-2017

Schulz, R., Drayer, R.A., Rollman, B.L., 2002. Depression as a risk factor for non-suicide mortality in the elderly. Biol. Psychiatry 52, 205–25. https://doi.org/10.1016/S0006-3223(02)01423-3

Shah, A.A., Han, J.Y., 2015. Anxiety. Continuum (Minneap. Minn). 21, 772–82. https://doi.org/10.1212/01.CON.0000466665.12779.dc

Simon, G.E., Rossom, R.C., Beck, A., Waitzfelder, B.E., Coleman, K.J., Stewart, C., Operskalski, B., Penfold, R.B., Shortreed, S.M., 2015. Antidepressants Are Not Overprescribed for Mild Depression. J. Clin. Psychiatry 8, 1627–1632. https://doi.org/10.4088/JCP.14m09162

Thomas, M., Price, D., 2008. Impact of comorbidities on asthma. Expert Rev. Clin. Immunol. 4, 731–742. https://doi.org/10.1586/1744666X.4.6.731

Thompson, A., Hunt, C., Issakidis, C., 2004. Why wait? Reasons for delay and prompts to seek help for mental health problems in an Australian clinical sample. Soc. Psychiatry Psychiatr. Epidemiol. 39, 810–7.

U.S. Preventive Services Task Force, 2009. Screening for depression in adults: U.S. preventive services task force recommendation statement. Ann. Intern. Med. 151, 784–92. https://doi.org/10.7326/0003-4819-151-11-200912010-00006

Verger, P., Saliba, B., Rouillon, F., Kovess-Masféty, V., Villani, P., Bouvenot, G., Lovell, A., 2008. Determinants of coprescription of anxiolytics with antidepressants in general practice. Can. J. Psychiatry 53, 94–103. https://doi.org/10.1177/070674370805300204

Wang, P.S., Berglund, P.A., Olfson, M., Pincus, H.A., Wells, K.B., Kessler, R.C., 2005. Failure and delay in initial treatment contact after first onset of mental disorders in the National Comorbidity Survey Replication. Arch. Gen. Psychiatry 62, 603–613. https://doi.org/10.1001/archpsyc.62.6.603

Weitz, E., Hollon, S.D., Kerkhof, A., Cuijpers, P., 2014. Do depression treatments reduce suicidal ideation? The effects of CBT, IPT, pharmacotherapy, and placebo on suicidality. J. Affect. Disord. 167, 98–103. https://doi.org/10.1016/j.jad.2014.05.036

Wen, X.J., Wang, L.M., Liu, Z.L., Huang, A., Liu, Y.Y., Hu, J.Y., 2014. Meta-analysis on the efficacy and tolerability of the augmentation of antidepressants with atypical antipsychotics in patients with major depressive disorder. Braz. J. Med. Biol. Res. 47, 605–16. https://doi.org/10.1590/1414-431x20143672

Wong, J., Motulsky, A., Abrahamowicz, M., Eguale, T., Buckeridge, D.L., Tamblyn, R., 2017. Off-label indications for antidepressants in primary care: descriptive study of prescriptions from an indication based electronic prescribing system. BMJ 356, j603. https://doi.org/10.1136/bmj.j603

Wray, N.R., Ripke, S., Mattheisen, M., Trzaskowski, M., Byrne, E.M., Abdellaoui, A., Adams, M.J., Agerbo, E., Air, T.M., Andlauer, T.M.F., Bacanu, S.-A., Bækvad-Hansen, M., Beekman, A.F.T., Bigdeli, T.B., Binder, E.B., Blackwood, D.R.H., Bryois, J., Buttenschøn, H.N., Bybjerg-Grauholm, J., Cai, N., Castelao, E., Christensen, J.H., Clarke, T.-K., Coleman, J.I.R., Colodro-Conde, L., Couvy-Duchesne, B., Craddock, N., Crawford, G.E., Crowley, C.A., Dashti, H.S., Davies, G., Deary, I.J., Degenhardt, F., Derks, E.M., Direk, N., Dolan, C. V., Dunn, E.C., Eley, T.C., Eriksson, N., Escott-Price, V., Kiadeh, F.H.F., Finucane, H.K., Forstner, A.J., Frank, J., Gaspar, H.A., Gill, M., Giusti-Rodríguez, P., Goes, F.S., Gordon, S.D., Grove, J., Hall, L.S., Hannon, E., Hansen, C.S., Hansen, T.F., Herms, S., Hickie, I.B., Hoffmann, P., Homuth, G., Horn, C., Hottenga, J.-J., Hougaard, D.M., Hu, M., Hyde, C.L., Ising, M., Jansen, R., Jin, F., Jorgenson, E., Knowles, J.A., Kohane, I.S., Kraft, J., Kretzschmar, W.W., Krogh, J., Kutalik, Z., Lane, J.M., Li, Y.Y., Li, Y.Y., Lind, P.A., Liu, X., Lu, L., MacIntyre, D.J., MacKinnon, D.F., Maier, R.M., Maier, W., Marchini, J., Mbarek, H., McGrath, P., McGuffin, P., Medland, S.E., Mehta, D., Middeldorp, C.M., Mihailov, E., Milaneschi, Y., Milani, L., Mill, J., Mondimore, F.M., Montgomery, G.W., Mostafavi, S., Mullins, N., Nauck, M., Ng, B., Nivard, M.G., Nyholt, D.R., O’Reilly, P.F., Oskarsson, H., Owen, M.J., Painter, J.N., Pedersen, C.B., Pedersen, M.G., Peterson, R.E., Pettersson, E., Peyrot, W.J., Pistis, G., Posthuma, D., Purcell, S.M., Quiroz, J.A., Qvist, P., Rice, J.P., Riley, B.P., Rivera, M., Saeed Mirza, S., Saxena, R., Schoevers, R., Schulte, E.C., Shen, L., Shi, J., Shyn, S.I., Sigurdsson, E., Sinnamon, G.B.C., Smit, J.H., Smith, D.J., Stefansson, H., Steinberg, S., Stockmeier, C.A., Streit, F., Strohmaier, J., Tansey, K.E., Teismann, H., Teumer, A., Thompson, W., Thomson, P.A., Thorgeirsson, T.E., Tian, C., Traylor, M., Treutlein, J., Trubetskoy, V., Uitterlinden, A.G., Umbricht, D., Van der Auwera, S., van Hemert, A.M., Viktorin, A., Visscher, P.M., Wang, Y., Webb, B.T., Weinsheimer, S.M., Wellmann, J., Willemsen, G., Witt, S.H., Wu, Y., Xi, H.S., Yang, J., Zhang, F., Arolt, V., Baune, B.T., Berger, K., Boomsma, D.I., Cichon, S., Dannlowski, U., de Geus, E.C.J., DePaulo, J.R., Domenici, E., Domschke, K., Esko, T., Grabe, H.J., Hamilton, S.P., Hayward, C., Heath, A.C., Hinds, D.A., Kendler, K.S., Kloiber, S., Lewis, G., Li, Q.S., Lucae, S., Madden, P.F.A., Magnusson, P.K., Martin, N.G., McIntosh, A.M., Metspalu, A., Mors, O., Mortensen, P.B., Müller-Myhsok, B., Nordentoft, M., Nöthen, M.M., O’Donovan, M.C., Paciga, S.A., Pedersen, N.L., Penninx, B.W.J.H., Perlis, R.H., Porteous, D.J., Potash, J.B., Preisig, M., Rietschel, M., Schaefer, C., Schulze, T.G., Smoller, J.W., Stefansson, K., Tiemeier, H., Uher, R., Völzke, H., Weissman, M.M., Werge, T., Winslow, A.R., Lewis, C.M., Levinson, D.F., Breen, G., Børglum, A.D., Sullivan, P.F., 2018. Genome-wide association analyses identify 44 risk variants and refine the genetic architecture of major depression. Nat. Genet. 50, 668–681. https://doi.org/10.1038/s41588-018-0090-3

Wulsin, L.R., Vaillant, G.E., Wells, V.E., 1999. A systematic review of the mortality of depression. Psychosom. Med. 61, 6–17.

Yan, X.-Y., Huang, S.-M., Huang, C.-Q., Wu, W.-H., Qin, Y., 2011. Marital Status and Risk for Late Life Depression: A Meta-Analysis of the Published Literature. J. Int. Med. Res. 39, 1142–1154. https://doi.org/10.1177/147323001103900402

Young, B.A., Von Korff, M., Heckbert, S.R., Ludman, E.J., Rutter, C., Lin, E.H.B., Ciechanowski, P.S., Oliver, M., Williams, L., Himmelfarb, J., Katon, W.J., 2010. Association of major depression and mortality in Stage 5 diabetic chronic kidney disease. Gen. Hosp. Psychiatry 32, 119–124. https://doi.org/10.1016/j.genhosppsych.2009.11.018

Zhou, L., Baughman, A.W., Lei, V.J., Lai, K.H., Navathe, A.S., Chang, F., Sordo, M., Topaz, M., Zhong, F., Murrali, M., Navathe, S., Rocha, R.A., 2015. Identifying Patients with Depression Using Free-text Clinical Documents. Stud. Health Technol. Inform. 216, 629–633. https://doi.org/10.3233/978-1-61499-564-7-629

